# FeS-cluster coordination of vertebrate thioredoxin regulates suppression of hypoxia-induced factor 2α through iron regulatory protein 1

**DOI:** 10.1101/2020.08.04.235721

**Authors:** Carsten Berndt, Eva-Maria Hanschmann, Laura Magdalena Jordt, Manuela Gellert, Leonie Thewes, Clara Ortegón Salas, Gereon Poschmann, Christina Sophia Müller, Yana Bodnar, Susanne Schipper, Oliver Handorf, Ricardo Nowack, Jean-Marc Moulis, Carola Schulzke, Volker Schünemann, Christopher Horst Lillig

## Abstract

Iron-regulatory protein 1 (IRP1), a central regulator of iron metabolism in vertebrates, also affects cellular response to hypoxia. IRP1 binds to the iron-responsive element (IRE) in the mRNA encoding hypoxia-inducible factor (HIF) 2α, thereby blocking the translation of the HIF2α-mRNA, and allowing the transcriptional regulation of, e.g., erythropoiesis. Here, we characterize the oxidoreductase thioredoxin 1 (Trx1) as a new regulator of hypoxia signaling. Human and murine Trx1 complex iron-sulfur clusters using one of the active site cysteinyl residues and a vertebrate-specific additional cysteinyl residue outside the active site. FeS-Trx1 is inactive, activated apo-Trx1 reduces cysteinyl residues in the binding pocket of IRP1/apo-Aconitase 1, which allows IRP1 to bind IREs in regulated mRNAs. Therefore, translation of the HIF2α mRNA requires either sufficient iron supply or the lack of reducing power of the Trx system under iron-limiting conditions. FeS-Trx1 thus links both redox and iron homeostasis to hypoxia responses.

## Introduction

Redox regulation, iron homeostasis, and the hypoxic response strongly depend on each other. Until now, however, mechanistic insights into these crosstalks are sparse. Thioredoxin (Trx) is a small oxidoreductase catalyzing thiol-disulfide exchange reactions with two cysteinyl residues in its CxxC active site motif (overview in ^1^). The dedicated flavo- and selenoenzyme Trx reductase (TrxR) reduces NADPH-dependent Trx. Mammalian cells contain two Trx/TrxRs, *i.e.*, cytosolic Trx1/TrxR1 and mitochondrial Trx2/TrxR2. Trxs were first described as electron donors for ribonucleotide reductase ^2^. Today, Trxs are better known as major regulators of various cellular functions through the redox regulation of thiol-disulfide switches and other redox modifications in key proteins ^3,4^.

Cellular functions, especially redox-controlled functions, depend on the presence of specific organic or inorganic cofactors. Iron is incorporated, *e.g.*, as heme or iron-sulfur (FeS) center into proteins. These proteins are essential to facilitate the transport of oxygen or electron transfer reactions, for instance, in the respiratory chain, DNA replication and repair, mRNA translation, and cellular metabolism ^5^. Iron deficiency can lead to anemia, iron overload can induce cellular toxicity, and is associated with various pathologies. Frequently, these can be traced back to the formation of hydroxyl radicals via the so-called Fenton reaction, see for instance ^4,6^. Because cellular iron levels need to be tightly controlled, organisms have developed effective mechanisms for iron regulation. Whereas lower organisms utilize transcriptional regulation to control iron homeostasis, vertebrates have developed a post-transcriptional mechanism via iron regulatory proteins (IRPs) ^7–9^ at the cellular level that complements systemic regulations. Activities of IRP1 and IRP2 are regulated in different ways. IRP1 is active under iron deficiency when its [4Fe-4S] cofactor is lost; the holo-protein functions as aconitase. Thus, the FeS cluster in IRP1 is considered to have an iron-sensing function ^8^. For RNA binding, cysteines of IRP1 need to be in a reduced state ^10^. The activity of IRP1 after nitrosylation, a cysteine modification, can be restored by Trx1 ^11^. IRP2, on the other hand, has not been demonstrated to bind a FeS cluster and is regulated via ubiquitination and proteasomal degradation. Ubiquitination depends on the F-box/LRR-repeat protein 5 (FBXL5), whose stability depends on a di-iron center ^12,13^. Moreover, binding between IRP2 and FBXL5 depends on the redox state of a FBXL5 coordinated [2Fe2S] cluster ^14^. IRPs 1 and 2 bind to specific mRNA structures, called iron-responsive elements (IREs), such as the transferrin receptor (TfR), required for iron uptake, the iron storage protein ferritin, and the iron export protein ferroportin. For approximately ten years, it has been known that also the 5’UTR of hypoxia-inducible factor 2α (HIF2α) mRNA contains an IRE ^15,16^. Although both IRPs can bind this IRE ^15^, *in vivo* experiments suggest that IRP1 dominates the suppression of HIF2α mRNA translation under iron-limiting conditions ^15–19^.

HIF2α is a critical part of the hypoxic response signaling pathway. In the presence of oxygen, the alpha subunits of all hypoxia-inducible factors are hydroxylated at conserved proline residues by specific HIF prolyl-hydroxylases. These hydroxylated forms are flagged for proteasomal degradation by the von-Hippel-Lindau E3 ubiquitin ligase ^20^. In the absence of oxygen, *i.e.*, hypoxia, hydroxylation fails, and the proteins translocate into the nucleus, acting as transcription factors. HIF2α controls erythropoiesis ^21^ and the growth and progression of various forms of cancer ^22^.

Here, we show that human and murine cytosolic Trx1s coordinate a [2Fe-2S]^2+^ cluster and explain the structural changes allowing the coordination of the cofactor. The loss of this FeS center activates the thiol-disulfide oxidoreductase activity of the protein. Active Trx1 is required to reduce cysteinyl residues in apo-IRP1, a prerequisite for their binding to IREs. The suppression of HIF2α translation under hypoxic – but iron-limited – conditions depends critically on the reductive capacity of the cytosolic Trx1/TrxR1/NADPH system.

## Results

During regular purification of recombinant Trxs expressed in *E. coli*, we noted a distinct brown-yellowish color of the human and mouse Trx1 proteins.

### Mammalian Trx1s complex a [2Fe-2S]^2+^ cluster

The UV-Vis spectra of the recombinant human and mouse proteins freshly purified from *E. coli* cultures display two peaks at approx. 335 and 420 nm, in addition to the peak of the aromatic side chains at 280 nm (Fig. 1a). These absorption features are characteristic of FeS clusters. After purification, colorimetric analyses revealed 0.48 ± 0.14 iron and 0.45 ± 0.12 sulfides per hTrx1 monomer. The freshly purified protein was analyzed by gel filtration chromatography, separating it into a monomeric and a dimeric fraction (Fig. 1a inset, Fig. S1). From these, only the dimeric fraction (27.71 kDa, theoretical MW of hTrx1 dimer: 27.89 kDa) displayed the additional absorption features, whereas the monomeric fraction (13.95 kDa, theoretical MW of Trx1 monomer: 13.90 kDa) was completely devoid of them (Fig. 1a). Reduction of the recombinant holo-proteins by DTT led to the loss of all spectral features characteristic for FeS clusters.

**Figure 1.**
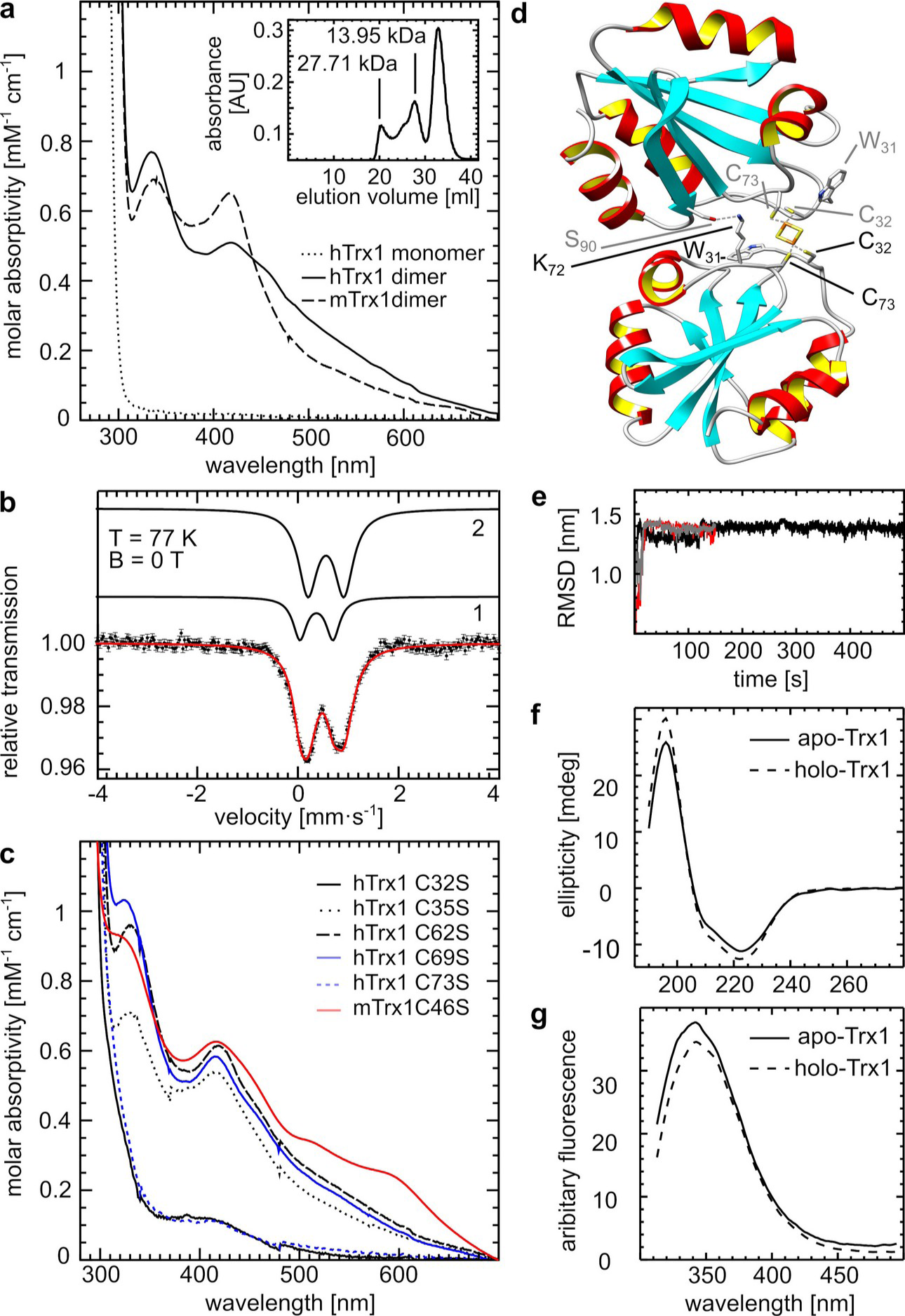
Identification of a [2Fe-2S]^2+^ cluster in human and mouse Trx1. a) UV-Vis spectroscopy of recombinantly expressed monomeric and dimeric human Trx1 and mouse Trx1 after purification. The spectra of the human Trx1 correspond to the fractions separated by gelfiltration shown in the inset. Inset: Chromatogram of the gelfiltration chromatography separating monomeric apo-Trx1 and dimeric holo-Trx1. b) Mössbauer spectra of reconstituted human Trx1 obtained at 77 K. The simulated spectra of fractions 1 and 2 (see text and table 1) were included on top of the recorded spectra. c) UV-Vis spectroscopy after purification of recombinant human and mouse Trx1 with individually mutated cysteinyl residues as indicated. d) Model of the holo-Trx1 complex after MD simulation (the most representative structure is shown). The FeS ligands and two amino acids stabilizing the dimer are highlighted. e) Root mean square deviations (RMSD) of the backbone of holo-Trx1 complex while three independent molecular dynamics simulations f-g) CD spectra (f) and fluorescence spectra at 294 nm excitation (Trp fluorescence, g) of monomeric apo-Trx1 and dimeric holo-Trx1 after gel filtration chromatography.

**Table 1.** Mössbauer parameters of reconstituted holo-Trx1. Holo-Trx1 was reconstituted *in vitro* and concentrated to 2.5 mM. The protein was frozen in liquid nitrogen and analyzed by Mössbauer spectroscopy at 77 K as outlined in the experimental procedures, see also Fig. 1.

| parameter | fraction 1 | fraction 2 |
| --- | --- | --- |
| *without external magnetic field* |  |  |
| $\delta$ (mm s<sup>-1</sup>) | 0.35 ± 0.02 | 0.56 ± 0.02 |
| $\Delta E_Q$ (mm s<sup>-1</sup>) | 0.66 ± 0.03 | 0.72 ± 0.03 |
| $\Gamma$ (mm s<sup>-1</sup>) | 0.33 ± 0.03 | 0.43 ± 0.03 |
| area (%) | 22 ± 2 | 78 ± 2 |

An ideal method for identifying and characterizing iron centers is Mössbauer spectroscopy. Hence, apo-human Trx1 was subjected to FeS cluster reconstitution to enrich the holo complexes with ^57^Fe. Mössbauer spectra were recorded for holo-Trx1 at 77 K (Fig. 1b) and 4.6 K in external magnetic fields (Fig. S2a-b). The spectra were simulated with two components (fractions 1 and 2) with a relative contribution of ca. 20 % and 80 %, respectively (Fig. 1b, Fig. S2a-b, Tab. 1, Tab. S1). Fraction 1 shows an isomer shift of δ = 0.35 mm s^-1^ and a quadrupole splitting of ΔE_Q_ = 0.66 mm s^-1^. δ is slightly higher than the values reported for high spin Fe^3+^ (S = 5/2) in a tetrahedral sulfur environment (δ = 0.20 - 0.32 mm s^-1^) ^22^. It is important to note that both δ and ΔE_Q_ are significantly smaller than those reported for [4Fe-4S]^2+^ clusters ^23^. The simulations of the spectra recorded in an external field of 0.1 T (Fig. S2A) and 5 T (Fig. S2b) are based on a diamagnetic species for fraction 1. Hence we conclude that this fraction represents a diamagnetic [2Fe-2S] ^2+^ cluster in which the antiferromagnetic coupling of the two paramagnetic Fe^3+^ ions results in an S = 0 state ^23–25^. Fraction 2 is characterized by δ = 0.56 mm s^-1^ and ΔE_Q_ = 0.72 mm s^-1^. These Mössbauer parameters are characteristic for high spin Fe^3+^ with octahedral nitrogen or oxygen coordination. The broad magnetic pattern observed at 4.6 K and fields of 0.1 and 5 T indicate the presence of small super-paramagnetic clusters that could originate from side reactions during the reconstitution process, possibly also from the decomposition of [2Fe-2S]^2+^ clusters ^26,27^.

The Mössbauer spectra of fraction 1 demonstrate the presence of [2Fe-2S]^2+^ clusters in dimeric holo-Trx1. Reconstitution of the FeS cluster in apo-Trx1 revealed approximately 30 ± 14 % holo-Trx1 (Mössbauer and UV-Vis spectroscopy), 40 ± 16 % of Trx1 were in the holo-form directly after purification from *E. coli* (UV-Vis spectroscopy). To identify the residues responsible for the coordination of the cluster, we individually mutated each of the five Cys residues of human Trx1 to Ser. To exclude that a loss of cluster ligation in any of the mutants was caused by the structural instability of the proteins, we confirmed their thermal stability by differential scanning calorimetry or fluorimetry (Fig. S3a, Tab. S2). Only the loss of one of two Cys residues led to the loss of the cluster binding, *i.e.* the more N-terminal active site residue C32 and the extra C73 outside the active site (Fig. 1c). Mouse Trx1 contains an additional C46, however, a C46S mutant was still able to coordinate the FeS cluster (Fig. 1c).

Can a [2Fe-2S]^2+^ cluster be complexed at the dimeric interface of two Trx molecules by a pair of C32/C73 ligands, as simulated in Fig. 1d? In various structures of human Trx1 deposited in the protein data bank, the sulfur groups of these two residues are separated by a distance of approx. 10 Å (Fig. S4a). This distance is too far for the joint coordination of the cofactor. However, as demonstrated by molecular modeling, only a small conformational change of the loop connecting helix 3 and strand 4 may bring the two sulfur atoms in a perfect distance for the coordination of the cluster, *i.e.,* 3.4 Å (Fig. S4a). The molecular dynamics (MD) simulations of the proposed model show no significant change in the overall secondary structure after the first 100 ns of the simulation (Fig.1e), thus suggesting the stability of the proposed model. A slight conformational difference between holo- and apo-Trx1 was confirmed by CD spectroscopy (Fig. 1f). These changes are proposed to go along with conformational changes of the two W31 residues (Fig. 1d), which should alter the local environment of the indole ring systems. Formation of the holo complexes induces quenching of the fluorescence of these (only) tryptophanyl residues (Fig. 1g). The solvatochromic nature of the Trp residues makes it sensitive to changes in the polarity of the local environment ^28^. Quenching can arise from changes in the interaction with the solvent water molecules and altered interactions with other side chains in the complex. The conformational changes of the loop structure proposed to accommodate [FeS] binding also affect the orientation of lysyl residue K72. Our model suggests that the dimeric holo complex may be stabilized by a direct interaction between the ε-amino groups of the K72 residues with the carboxyl groups of S90 of the respective other subunit (Fig. 1d). In line with this hypothesis, Trx1K72E lost the spectral characteristics of the [2Fe-2S]^2+^ cluster coordinated by wt Trx1 (Fig. S4b) and is less stable (Fig. S3b, Tab. S2). However, the redox activity was only affected to a minor degree since the mutated form could still reduce insulin disulfides and can be reduced by TrxR1. The *K_m_* of TrxR1 towards Trx1K72E increased from 3.0 ± 0.2 to 4.4 ± 1.7 µmol·l^-1^, the *v_max_* slightly increased from 2.7 ± 0.1 to 3.1 ± 0.5 s^-1^ (Fig. S4c). The decreased affinity of TrxR for Trx1K72E was likely the result of changes in the electrostatic properties of the mutant (Fig. S4d-e). In addition to K72, several other amino acids might contribute to the stabilization of the dimer (Fig. S4f). All these data confirm the formation of a holo-dimer using cysteinyl residues C32 and C73 as FeS cluster ligands. C73 is present neither in the mammalian mitochondrial Trx2 nor in bacterial Trxs, and none of these proteins contain a chromophore (Fig. S5).

Does human Trx1 bind the iron cofactor also *in vivo* as suggested by incorporation of the FeS clusters into Trx1 during recombinant expression in *E. coli* (Fig. 1a)? To answer this question, we propagated Jurkat cells of human origin in the presence of transferrin-bound radioactive ^55^Fe. Trx1 and, as a control, glutaredoxin (Grx) 1, which does not bind iron ^29,30^, were immuno-precipitated from the cell extract and analyzed by scintillation counting for iron binding. Significant amounts of iron co-precipitated with Trx1 but not Grx1 (Fig. 2a).

**Figure 2.**
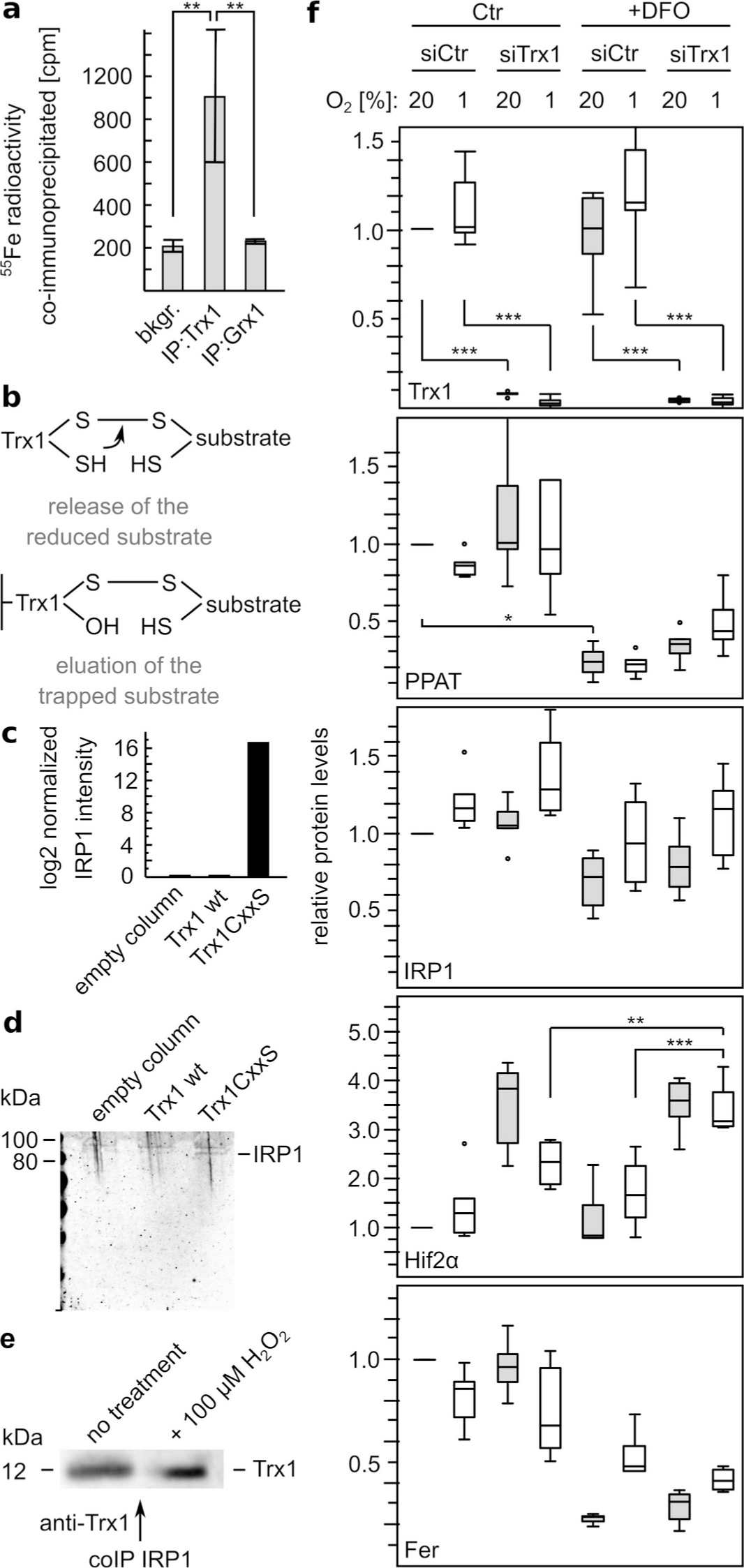
Trx1 interacts with IRP1 and affects the response on iron-depletion and hypoxia. a) Co-immunoprecipitation of ^55^Fe with Trx1 from Jurkat cell extracts propagated with transferrin-bound ^55^Fe as main iron source (n=3, mean ± SD). b) Schematic mechanism of substrate trapping by a column coupled Trx1 mutant. c-d) Identification of IRP1 as substrate of Trx1 using HeLa cell extracts. c) Quantification of IRP1 peptides via mass spectrometry in the eluted interaction partners. d) Identification of IRP1 as substrate after Western blotting of the eluted trapped substrates. e) Co-immunoprecipitation using anti-IRP1 antibodies revealed Trx1 as interacting protein in Jurkat cells ± H_2_O_2_ treatment after Western blotting. f) Analysis of the protein levels of Trx1, phosphoribosylpyrophosphate amidotransferase (PPAT), iron regulatory protein 1 (IRP1), hypoxia-induced factor 2α (HIF2α), and ferritin (Fer) in control cells (siCtr) and cells depleted of Trx1 by siRNA treatment (siTrx1). These cells were cultivated with or without the iron chelator deferoxamine (DFO) under 20 % or 1 % oxygen to induce both, iron-limiting and hypoxic conditions. The box plots depict the medians as center lines; box limits indicate the 25^th^ and 75^th^ percentiles as determined by R software; whiskers extend 1.5 times the interquartile range from the 25^th^ and 75^th^ percentiles, outliers are represented by dots (n = 6, *: p<0.05, **: p<0.01, ***: p<0.001, Students T-test).

### Trx1 and cellular iron metabolism

Since FeS cluster coordination blocks the CxxC active site, we assumed a Trx1-dependent connection between redox and iron regulation. Using Trx1C35S, a mutant lacking the second resolving cysteinyl residue of the active site, we were able to trap and identify potential substrates of Trx1 in HeLa cells (Fig. 2b). IRP1, the post-transcriptional iron regulator located in the cytosol of mammalian cells, was one of these substrates (Fig. 2c-d). In addition, Trx1 was confirmed *vice versa* as an interaction partner of IRP1 using co-immunoprecipitation under untreated and H_2_O_2_-treated conditions in Jurkat cells (Fig. 2e). Based on this, we analyzed the consequence of Trx1 depletion on the level of IRP1-regulated proteins. Since IRP1 is also an essential regulator of hypoxia response, HeLa cells were cultivated at 1 % oxygen to inhibit the proteasomal degradation of HIF2α and treated with the iron-chelator deferoxamine (DFO), thus provoking both iron-depletion and hypoxia responses. Treatment with specific siRNA lowered Trx1 levels to less than 10 % (Fig. 2f). Significant lower levels of phosphoribosylpyrophosphate amidotransferase (PPAT), a metabolic enzyme that is degraded when the incorporation of its FeS cluster fails, showed effective iron depletion (Fig. 2f). However, depletion of Trx1 had no relevant effect on the diminished cytosolic FeS cluster biogenesis. The cellular amount of IRP1 itself was almost unaffected (Fig. 2f). It was slightly lower in iron-depleted cells and, under all tested conditions, a bit higher in Trx1-lacking cells (132 ± 9 %). The HIF2α encoding mRNA, one of the main targets of IRP1, contains an IRE in its 5’ region. Hence, the translation of HIF2α is blocked by IRP1 binding. This block is not present in cells lacking Trx1 (Fig. 2f). In these cells, the HIF2α levels increased to 231 ± 49 % and 342 ± 57 % if iron was absent. Surprisingly, levels of the iron storage protein ferritin did not rise upon Trx1 deficiency (Fig. 2f), although the respective mRNA contains an IRE in its 5° region.

### Human Trx1 reduces Iron-Regulatory Protein 1 in vitro and in vivo

Next, we tested recombinant oxidized human apo-IRP1 as a substrate for Trx1 in a coupled assay with TrxR1 following NADPH consumption. As depicted in Fig. 3a, apo-Trx1 was able to reduce and thus activate IRP1 *in vitro* with a K_m_ of 18.5 µM and a v_max_ of 0.41 s^-1^. To confirm that only apo-IRP1 is a substrate, we performed the described assay in an anaerobic chamber using purified holo-IRP1 with and without cluster removal. With 16 µM apo-IRP1, we measured an enzymatic Trx1 activity of 5.94 ± 2.82 µM/min, and with holo-IRP1 3.12 ± 2.4 µM/min (Fig. 3b), indicating that apo-IRP1 is the main target for Trx1. In line, aconitase (holo-IRP1) activity in the cytosol of hypoxic HeLa cells was not affected by Trx1 depletion (Fig. S6).

**Figure 3.**
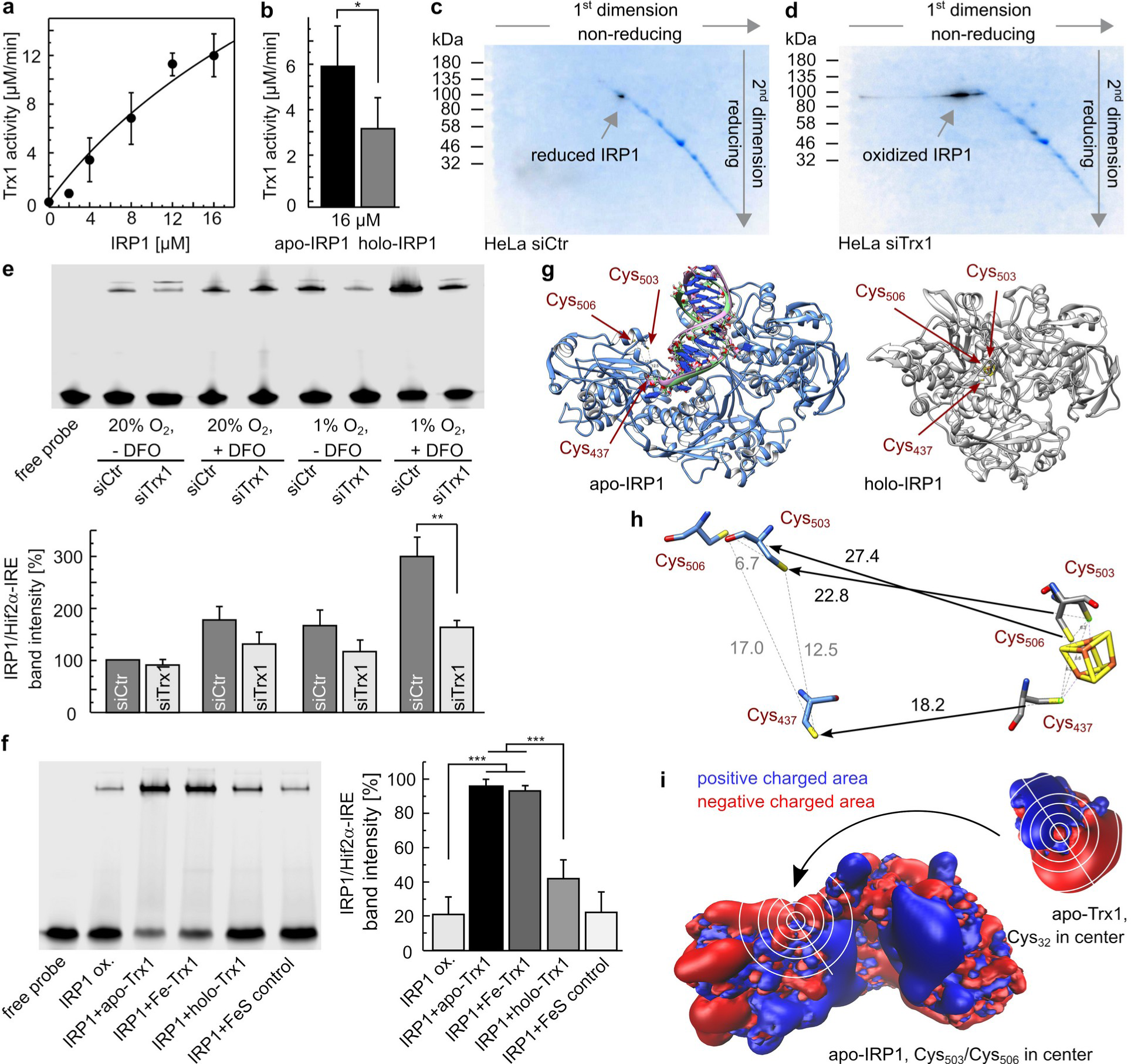
Trx1 reduces IRP1 and regulates IRP1 binding to Hif2α-IRE. a) Reduction of different concentrations of recombinant IRP1 as indicated by Trx1 *in vitro* in a coupled kinetic assay with rat TrxR1 and NADPH measuring NADPH consumption at 340 nm. The solid line is a non-linear curve fitting of the data points (n = 4, mean ± SD) to the Michaelis-Menten equation. b) Using the same assay as in a), holo- and apo-IRP1 were used as substrates. The assay was performed under anaerobic conditions (n = 3, mean ± SD). c-d) Diagonal 2-dimensional SDS PAGE under non-reducing (first dimension) and reducing conditions (second dimension) reveals the redox state of IRP1 in control (c) and Trx1-depleted HeLa cells (d) cultivated at 1% O_2_ in the presence of deferoxamine (DFO). The pictures show overlays of the total amount of proteins in blue (Coomassie staining) and a Western blot staining of IRP1 in black. e) Electrophoretic mobility shift assays (EMSAs) with extracts of HeLa cells cultivated under 20 % or 1 % O_2_, ± iron chelator deferoxamine (DFO), and ± Trx1 depletion (siTrx1 or siCtr). Extracts were mixed with infrared-labeled Hif2α-IRE probe to determine RNA binding capacity of IRP1 (n = 6-14, mean ± SEM, **: p<0.001, Students T-test). f) Recombinant IRP1 was incubated with apo-Trx, holo-Trx (after *in vitro* reconstitution), Trx-Fe (Trx1 incubated with (NH_4_)_2_Fe(SO_4_)_2_), and FeS control mixture (*in vitro* reconstitution mixture without Trx1) in presence of a labeled Hif2α-IRE probe to perform an EMSA Quantification shows mean ± SEM (n = 3-11, **: p<0.01, ***: p<0.001, Students T-test). g) Model structure of apo-IRP1 with bound IREs (left blue structure based on pdb codes 3sn2 and 3snp) and holo-IRP1/Aconitase (right grey structure based on pdb code 2b3x). h) Orientation, distances, and movements (arrows) of cysteinyl residues 437, 503, and 506 during apo to holo transition. i) Electrostatic isosurfaces of apo-IRP1 and hTrx1 at ± 25 mV display complementary binding surfaces.

Does Trx1 also fulfill this function *in vivo*? We analyzed the redox state of IRP1 in Trx1-depleted HeLa cells compared to cells treated with control siRNA via two-dimensional diagonal gel electrophoresis. In the first dimension, the proteins were separated by non-reducing SDS PAGE; the second dimension was SDS PAGE under reducing conditions. In this assay, reduced proteins end up on a diagonal according to their molecular weight (Fig. 3c-d). Proteins with oxidized cysteinyl residues, however, drop below the diagonal. As demonstrated by an overlay of Coomassie-stained gel and Western blot detecting IRP1, all IRP1 in control cells was reduced (Fig. 3c). Upon Trx1-depletion, however, most of the IRP1 protein was oxidized (Fig. 3d). Trx1 is required for the reduction and thus activation of IRP1 *in vivo*.

### Trx1 regulates IRP1 binding to the Hif2α-IRE

The HIF2α mRNA has been demonstrated to be one of the major targets of IRP1 ^19^. It has long been known from *in vitro* assays that the binding of IRP1 to IREs requires the reduction of the cysteinyl residues in the IRE binding site, see *e.g.* ^10^. These residues also coordinate the FeS cluster within IRP1 that is lost upon iron depletion. These cysteinyl residues are prone to be oxidized once the cluster is gone. Therefore, we tested whether the reduction of IRP1 by Trx1 affects IRE binding. Whereas Trx1 depletion via siRNA (Fig. S7a) showed no impact on IRP1/HIF2α-IRE binding in whole lysates of cells cultivated under standard conditions, the increased IRP1/HIF2α-IRE binding in wildtype cells under iron-limited and hypoxic conditions (300 ± 36 % compared to untainted cells) was severely and significantly diminished in Trx1-depleted HeLa cell lysates (only 160 ± 17 %) in electrophoretic mobility shift assays (EMSAs, Fig. 3e). The diminished IRE binding capacity was slightly more pronounced in the purified cytosolic fraction (48 ± 15 %, Fig. S7b) compared to the whole lysate (57 ± 7 %, Fig. 3e). Moreover, decreased IRE binding upon Trx1 depletion was determined in EMSAs under hypoxic or iron depleting conditions alone in whole lysate (Fig. 3e) and even under standard conditions in the isolated cytosolic fraction (Fig. S7b). As shown before (Fig. 2f), the lack of Trx1 does not affect IRP1 levels, and activating oxidized IRP1 by the reduction of lysates with DTT further showed that decreased IRP1/HIF2α-IRE binding upon Trx1 depletion is not based on decreased levels of IRP1: the formation of IRP-IRE complexes increased in lysates of control cells after incubation with DTT only to 115 ± 16 %, but to 194 ± 56 % in lysates of Trx1 lacking cells (Fig. S7c). The formation of supershift complexes after the addition of anti-IRP1-antibodies confirmed IRP1-IRE probe complexes (Fig. S7d-e). EMSAs with purified recombinant IRP1 and Trx1 demonstrated direct reduction and activation of IRP1 by Trx1. 10 µM of Trx1 is as efficient as 10 mM β-mercaptoethanol in activating 1 µM IRP1, IRE binding increased to 359 ± 92 % in the presence of apo-Trx1 and 251 ± 51 % in the presence of β-mercaptoethanol (Fig. S7e). The blocking of the CxxC active site by coordination of the FeS cluster after reconstitution diminished the ability of Trx1 to activate IRP1 to 44 ± 12 %, whereas incubation with Fe^2+^, most likely leading to unspecific iron binding did not affect Trx1 activity in modulating IRP1/HIF2α-IRE binding (97 ± 4 %, Fig. 3f).

IRP1 contains nine cysteines. The three cysteinyl residues essential for both FeS cluster coordination in holo-IRP1 and IRE binding in apo-IRP1 are C437, C503, and C506 (Fig. 3g). These cysteines are close together in the holo-form (approximately 6 Å) but shift 18.2, 22.8, and 27.4 Å, respectively, to their positions in the apo-form where the distance between C437 and C503 or C506 is 12.5 and 17 Å (Fig. 3h). Based on the electrostatic surface potential maps of apo-IRP1 and apo-Trx1, we predict that the active site of Trx1 most likely attacks either C503 or C506 (Fig. 3i). Therefore, we hypothesize the formation of a disulfide in IRP1 between cysteinyl residues 437 and 503/506 after disassembly of the FeS cluster that inhibits IRE binding and the need for active Trx1 to reduce this disulfide and to rescue the IRE binding activity of apo-IRP1.

## Discussion

The present study characterized the human and murine cytosolic Trx1 as new FeS-proteins. The [2Fe-2S]^2+^ cluster is coordinated at the dimeric interface of two Trx molecules. Approximately a decade ago, we characterized human Grx2 as the first Trx family member containing an FeS cofactor ^30^. The [2Fe-2S]^2+^ cluster in Grx2 is coordinated by the N-terminal active site Cys of two monomers plus two additional non-covalently bound glutathione molecules ^31,32^. Because the glutathione used as FeS ligand is in permanent exchange with the free pool of glutathione, the iron complex was described as a redox sensor that could regulate the oxidoreductase activity depending on the redox state of the free glutathione pool ^31^. Holo-Grx2 is inactive as an oxidoreductase but appears to be involved in nitric oxide detoxification during neuroinflammation ^33^. Enzymatically active apo-Grx2 is required, for instance, for brain and vascular development ^34,35^ and promotes cell migration of cancer cells ^35,36^. The same mode of cluster coordination was also demonstrated for the human monothiol Grxs 3 and 5 ^29,37^ that function in iron homeostasis and FeS cluster biosynthesis, as well as for various Grxs from other species, for overviews; see, for instance ^38,39^. Fig. 4a depicts the overlay of the iron-coordinating thiol groups in human Trx1, Grx2, and Grx5, centered on the common N-terminal active site cysteinyl residue (C32 in Trx1), indicating that FeS cluster coordination by Trx family proteins needs a second thiol group in the suitable distance range to the active site thiol. Whereas Grxs insert this thiol via the bound glutathione, human Trx1 uses the side-chain of the not universally conserved C73 as the second thiol. The position of this second thiol appears to be variable due to the dimeric nature of the holo*-*complexes. The slightly shifted positions of the glutathione thiols and FeS centers are caused by alternative loop structures preceding the active sites. These differences are also the structural basis for the functional difference (oxidoreductases or FeS cluster transferases) between the two classes of vertebrate glutaredoxins ^40^. The different positions of the glutathione thiol cause a rotation of the two Grx monomers in the Grx2 and Grx5 holo-complexes relative to each other ^32,37^. This is likely also true for the Trx1 holo-complex, although Mössbauer spectroscopy revealed a slightly higher value of δ than usually found for [2Fe-2S] clusters coordinated by four sulfhydryl groups. This increase could be related to the presence of a nitrogen or an oxygen ligand ^24,25^. Recently, for the bishistidinyl-coordinated non-Rieske [2Fe-2S]^2+^ proteins, a value of δ = 0.36 mm s^-1^ has been reported for the bis-histidyl coordinated Fe^3+^ site at pH=6 ^41^. However, the only nitrogen or oxygen available for cluster coordination near the active site would be the histidyl residues introduced by the His-tag. Since we used both C- as well as N-terminal His-tags and were able to show that specific Trx1 mutants affect cluster coordination independent of the His-tag, we are confident that the cofactor is coordinated by cysteinyl residues 32 and 73. In retrospect, this coordination was already indicated by our work showing that only the additional removal of these two cysteinyl residues in human Trx1 inhibited [2Fe-2S] cluster coordination in a mutant lacking the *cis*-proline (P75 in human Trx1), another feature regulating FeS cluster coordination in members of the Trx family ^42^. However, recently, a [4Fe-4S] cluster coordinated by the residues of the two active site cysteinyl residues was described in *A. thaliana* Trx o ^43^. This Trx is located in mitochondria, contains a *cis*-prolyl residue, and lacks the additional C73, which serves as the second thiol in a similar position as in the FeS-Grxs. In addition, some Trx-type/Trx-related proteins were identified as FeS proteins, *e.g.* dimers of *E. granulosus* iron-sulfur Trx-related protein (IsTRP) and *T. brucei* Trx2 coordinate [2Fe 2S] clusters ^44,45^. Several mutants of *E. coli* Trx with engineered active site sequences are able to coordinate [2Fe-2S] clusters as well ^46,47^. In general, the Trx fold seems to be well suited for the coordination of metal cofactors, as shown by many other engineered and wildtype proteins ^48^. C73 is not a ubiquitously conserved residue. We have performed an extensive sequence comparison between Trxs from various species, classes, and kingdoms, see Fig. S8. This analysis identified the extra C73 as specific for vertebrate cytosolic Trxs, with only a few exceptions in arachnid species and one mollusk. C73 is not present in Trxs from bacteria, fungi, plants, insects, nematodes, or mitochondrial vertebrate Trxs. We have expressed and purified both *E. coli* Trx1 and human mitochondrial Trx2 in the same way as human and mouse Trx1 and did not detect any signs of cluster binding (Fig. S5). Hence, nearly all potential FeS-Trxs exist in the same evolutionary group that utilizes the IRP/IRE system for the regulation of iron uptake and storage (also included in Fig. S8); all of the C73-Trxs are cytosolic proteins. Only one previous study linked Trx1 to the IRP regulon. In macrophages, Trx1 was reportedly required to activate IRP1 after exposure to nitric oxide ^11^. Trx1 can reduce IRP1 i*n vitro* and *in vivo*, leading to increased IRE binding of HIF2α mRNA (Fig. 3). The importance of the reduction of the cysteinyl residues in the FeS cluster coordination site of IRP1, which is also the IRE binding site, has long been known from i*n vitro* assays, *e.g.* ^10^. Here, we focus on the redox state of cysteinyl residues. However, also other amino acids may contribute to the redox regulation of IRP1 activity, i.e. tyrosine nitration by peroxynitrite ^49^. When iron is not limiting, IRP1 (gene name ACO1) is a functional cytosolic aconitase with a [4Fe-4S] cluster in its active site that is ligated by three cysteinyl residues. This cluster is lost when iron becomes a limiting factor, IRP1 undergoes structural changes between holo- and apo-form, and the resulting apo-protein can bind IRE stem-loop structures in mRNAs (Fig. 3g). The three cysteinyl residues, C437, C503, and C506, bind the FeS cluster or the IRE. The redox state of C437 has been described before as important for IRE binding ^50^. Only mutation of this cysteinyl residue rescued the lost IRP1-IRE binding upon treatment with either N-ethyl-maleimide, a molecule blocking free thiols, or diamide, a compound that unspecifically induces the formation of disulfides. Of course, the formation of a disulfide needs two cysteinyl residues. The interaction between Trx and a substrate critically depends on complementary electrostatic properties ^51,52^. Based on these properties, Trx1 might reduce a disulfide between cysteinyl residues 437 and 503/506 (Fig. 3i). The formation of a disulfide that may prevent IRE-binding was suggested before ^53^. This is, however, in contrast to several reports describing the activation of IRP1 by H_2_O_2_, summarized in ^54^. Noteworthy, this activation can be observed only at low concentrations of H_2_O_2_ in cells and not in lysates. In intact cells, low levels of H_2_O_2_ most likely activate the Keap1/Nrf2 system, which leads to the expression of various enzymes counteracting oxidative damage, including Trx1, that subsequently promote the activation of IRP1. In line, recombinant holo-IRP1 lost its FeS cluster upon H_2_O_2_ treatment but did not gain IRE binding capacity ^55^.

**Figure 4.**
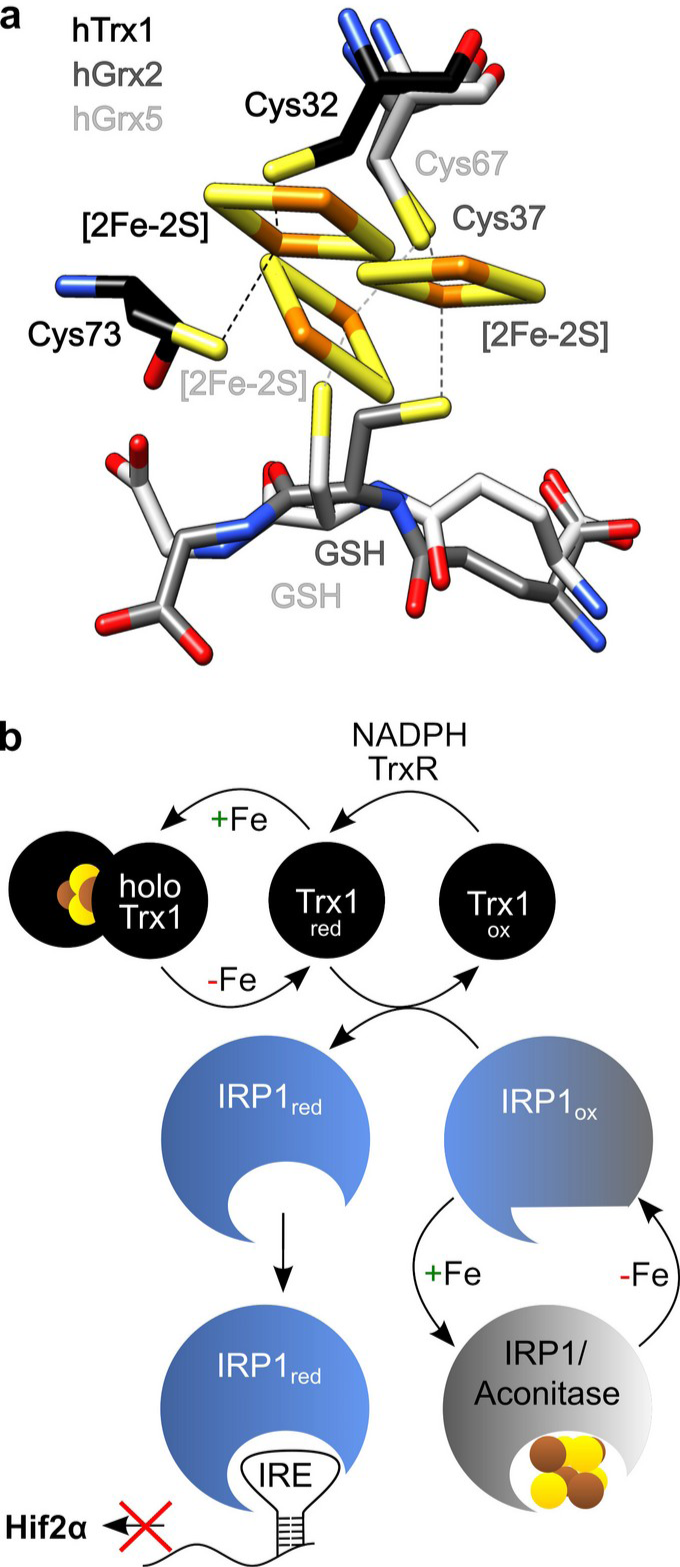
FeS cluster coordination in Trx family proteins. **a**) Alignment of the structures of human holo-Grx2 (pdb entry: 2ht9), holo-Grx5 (2wul), and holo-Trx1 (obtained by modeling in this study) depicting the position of the thiol groups recruited for binding of the [2Fe-2S] clusters. The FeS clusters bound to Grx2 and Grx5 were included in the image. b) Model for the activation of Trx1 and IRP1 by FeS cluster disassembly and redox regulation, for details see text.

Lack of Trx1 led to a massive increase in HIF2α, despite conditions of iron-limitation, while ferritin was hardly affected (Fig. 2f). That was unexpected as both RNAs contain the IRE at the 5’ region. However, as described before ^55^, a hypoxia-dependent increase in ferritin levels seems cell-specific. Moreover, ferritin iron-loading in the experiments of Fig. 2f may be different. Last, hypoxic and iron-scavenging kinetics might differ. In the case of cellular response to hypoxic conditions, a redox-dependent signaling element would be helpful. A comparison between the IREs of ferritin and HIF2α reveals two guanine bases absent in ferritin’s IRE. These guanine bases are located between C8 and the conserved CAGUG motif at the top of the IREs, the two areas forming the most bonds with IRP1 ^56^. Since guanine exists as an oxidized base changing RNA binding specificity ^57^, we do not want to exclude another level of regulation, *e.g.*, IRE oxidation ^49^. Mammalian IRP1 and IRP2 share functions ^59^ but are differently affected by hypoxia: IRP1 is inhibited, while IRP2 is activated ^60,61^, although data that suggest the activation of IRP1 during hypoxia have also been presented ^55,62^. Several studies, including analyses of knock-out mice, emphasized the pivotal function of IRP1 in suppressing HIF2α translation under iron-limiting conditions ^17–19^. HIF2α is an essential regulator of erythropoiesis, angiogenesis, cell proliferation, de-differentiation, and invasion of cancer cells via its various target proteins ^19,21,22,63^.

HIF2α is primarily regulated by suppressing its translation through IRP1 under iron-limiting conditions and the O_2_-dependent proteasomal degradation pathway. With redox regulation this study adds a third level of regulation to the IRP1-HIF2α signaling hub. The lack of Trx1 activity in cells leads to the loss of HIF2α suppression even under iron-limiting conditions. Fig. 4b summarizes this extended model of IRP1 activation and, thus, permission or suppression of HIF2α translation. Activation of both Trx1 and IRP1 requires conditions that lead to the loss of their iron-sulfur cofactors (Fig. 3e). Next, activation of IRP1 involves the reduction of the protein by Trx1 with electrons donated by TrxR1 and NADPH. The limitation of the reducing power of the Trx system may explain the limited ability of IRP to regulate exogenous IREs in transfected cellular systems ^64^. Only when both conditions are met, iron limitation and sufficient reducing power of the cytosolic Trx system, IRP1 can bind to the IRE in the 5’-end of the HIF2α mRNA and thus suppress its translation and accumulation of the transcription factor under hypoxic conditions. In support of this model, all mentioned HIF2α processes are also redox-regulated on multiple levels, many of which are dependent on Trx1, summarized in ^4^.

## Methods

### Cloning, protein expression and purification

Human and mouse Trx1 were cloned into the expression plasmids pet20b and pet15b as described before ^42,65^. Mutations were inserted by rolling circle mutagenesis using specific oligonucleotides (C32S: fwd: cagccacgtggtctgggccttg, rev: caaggcccagaccacgtggctg; C35S: fwd: tggtgtgggccttccaaaatgatc, rev: gatcattttggaaggcccacacca; C62S: fwd: gtggatgactctcaggatgttgc, rev: gcaacatcctgagagtcatccac; C69S: fwd: gttgcttcagagtctgaagtcaaatg, rev: catttgacttcagactctgaagcaac; C73S: fwd: gaagtcaaatccatgccaacattcc, rev: ggaatgttggcatggatttgacttc; K72E: fwd: tcagagtgtgaagtcgagtgcatgccaacattc, rev: gaatgttggcatgcactcgacttcacactctga). IRP1 was cloned by PCR from human cDNA using specific oligonucleotides (fwd: gctagcatgagcaacccattcgcac, rev: ctcgagcttggccatcttgcggatcatg), digested with NheI and XhoI, and cloned into the vector pET16b (Novagen). Proteins were produced in *E. coli* as His-tag proteins and were purified using the IMAC principle (GE healthcare). Following purification, proteins were buffered in PBS using NAP-5 columns (GE healthcare) or Zeba spin columns (Thermo Scientific). The thermal stability of Trx1 mutated forms was analyzed using differential scanning calorimetry or fuorimetry. 10 or 40 µM of the proteins in PBS were heated from 20 to 100 °C at 0.2 - 1 K per minute in a Micro Cal VP-DSC (Malvern Panalytical Ltd) and the heat flux was recorded or, for DSF, followed in a BioRad CFX qPCR instrument. The melting point (T_m_) of DSF was identified by fitting the melting curve with Boltzmann equation (eq.1). In this equation FU is the measured fluorescence signal from Sypro Orange.

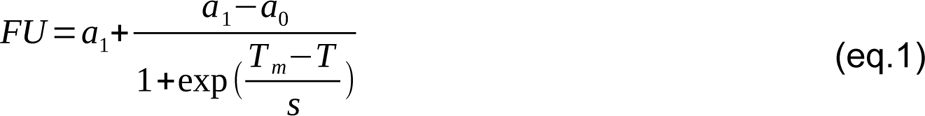

Gelfiltration chromatography for the separation of monomeric and dimeric Trx1 was performed as outlined in ^29^.

### In vitro reconstitution

Reconstitution of the FeS cluster was performed as described in ^31^ without GSH. For Mössbauer spectroscopy apo-Trx1 was reconstituted using (NH_4_)_2_ ^57^Fe(SO_4_)_2_ as iron source.

### Spectroscopy

Following purification, reconstitution, and buffering in PBS, proteins were desalted using NAP-5 columns, UV-Vis spectra were recorded using 100-250 µM of wildtype and mutant proteins with the Uvikon 922 photometer (Kontron Instruments) from 240-720 nm. Circular dichroism (CD) spectra were recorded with 10 µM apo- and holo-Trx1 in a 1 mm cuvette, scanning 0.2 nm steps, averaging 5 iterations, with the Jasco J-810 instrument. Tryptophanyl fluorescence was recorded with a Perkin Elmer LS50B fluoriometer (10 µmol·l^-1^ protein, excitation: 296 nm, 6 nm excitation slit and 4 nm emission slit).

A conventional Mössbauer spectrometer was operated in the constant acceleration mode in conjunction with a multichannel analyzer in the time-scale mode (WissEl GmbH) to record transmission Mössbauer spectra. The spectrometer was calibrated against α-iron at room temperature. The samples were cooled to 77 K with a flow cryostat (OptistatDN, Oxford Instruments). Field-dependent conventional Mössbauer spectra were obtained with a helium closed-cycle cryostat (CRYO Industries of America, Inc.) equipped with a superconducting magnet ^66^. The magnetic field direction parallel to the γ-ray beam. The spectral data were transferred from a multichannel analyzer to a PC for further analysis employing the public domain program Vinda ^67^ running on an Excel (2003) platform. Analysis of the spectra was performed by least-squares fits using Lorentzian line shapes with the linewidth Γ. Field-dependent spectra were simulated by means of the spin Hamilton formalism ^68^. Following *in vitro* reconstitution, human Trx1 was concentrated to 2.5 mM using the Stirred Ultrafiltration Cell (Millipore) and buffered in PBS using PD-10 columns (GE Healthcare). (NH_4_)_2_^57^Fe(SO_4_)_2_·6H_2_O was synthesized in a procedure adapted from ^69^. In brief, 102.9 mg (1.768 mmol, 1 eq) of ^57^Fe were dissolved in 2.6 ml of 1 molar sulfuric acid with continuous heating under N_2_ atmosphere. The resulting turquoise solution was filtered from the deposited coke. To this solution, 243.14 mg (1.84 mmol, 1 eq) ammonium sulfate in aqueous solution were added. Next, the solution was concentrated under reduced pressure and allowed to form a crystalline surface; the concentrated solution was heated and stored by 4 °C to allow crystallization. The resulting turquoise crystals were collected and the procedure was repeated for the remaining solution. In the end, 67.7 % of the initial iron was present as highly pure (NH_4_)_2_^57^Fe(SO_4_)_2_·6H_2_O crystals, as confirmed by X-ray analysis ^70^.

### Activity measurements

The activity of cytosolic aconitase was determined as described in ^71^. The activity of Trx1 in the reduction of IRP1 was measured in a coupled optical assay with rat thioredoxin reductase and NADPH as outlined in ^72^. The reduction of insulin in a coupled assay of Trx with TrxR has been performed following the reduction of NADPH spectrophotometrically. The assay mixture contained 50 mmol·l^-1^ Tris/HCl pH 7.5, 2 mmol·l^-1^ EDTA, 160 µmol·l^-1^ insulin (SIGMA-Aldrich), 150 µmol·l^-1^ NADPH, 1.25 nmol·l^-1^ recombinant rat TrxR1 (kind gift from Elias Arner, Stockholm), and Trx from 0 – 25 µmol·l^-1^. Holo-IRP1 used as substrate was purified under anaerobic conditions as described in ^73^. The FeS cluster was removed according to ^74^ by incubation of the protein with 200 µM H_2_O_2_ and 10 mM EDTA for 1 h at room temperature following filtration through a Zeba spin column (Thermo Scientific). The respective activity assays were performed in an anaerobic chamber (Coy Laboratory Products, USA).

### Cell culture

HeLa cells were cultured in low glucose (1g/l) DMEM medium, supplemented with 10 % FCS and 100 units/ml penicillin 0.1 mg/ml streptomycin at 37 °C in a humidified atmosphere containing 5 % CO_2_. Cells were transfected with specific siRNA against human Trx1 by electroporation. 3.5·10^6^ cells were resuspended in electroporation buffer (21 mM HEPES, 137 mM NaCl, 5 mM KCl, 0.7 mM Na_2_HPO_4_, 6 mM D-glucose, pH 7.15), were mixed with 15 µg of Trx1 siRNA (sense: guagauguggaugacuguc, antisense: gacagucauccacaucuac) or scrambled control siRNA (sense: cauucacucaggucauca, antisense: cugaugaccugagugaau) and were electroporated at 250 V, 1500 µF and 500 Ω using the BTX ECM 630. FCS was immediately added to the cells before seeding them in 1:5 conditioned medium. The following day the medium was changed. For sufficient knock-down, cells were transfected a second time after three days and cultivated for another 2 days at 20 % oxygen followed by 24 h at 1 % oxygen concentration. To reduce intracellular iron levels, cells were treated for 24 h with 100 µM deferoxamine (Abcam). Cells were harvested by trypsination, incubated with PBS containing 100 mM NEM for 5 min at RT. Cells were centrifuged and washed once in PBS, before 30 min lysis in 40 mM HEPES, 50 mM NaCl, 1 mM EDTA, 1 mM EGTA, 2 % CHAPS, complete protease inhibitor, and 100 mM NEM. Samples were stored at -80 °C. The knock-down efficiency, as well as the iron depletion was analyzed by Western blot using specific antibodies against Trx1 and iron-regulated proteins.

Jurkat cells were cultured in RPMI medium supplemented with 10 % FCS, 100 units/ml penicillin and streptomycin, and 2 mM glutamine at 37 °C in a humidified atmosphere containing 5 % CO_2_. Radiolabeling was done with 1 mCi, 2 µM transferrin-^55^Fe (Perkin-Elmer). Co-immunoprecipitation of ^55^Fe with Trx1 and Grx1 was performed as described for Grx3 in ^29^.

Cell fractionation was performed according to ^71^. Cells were resuspended in mitobuffer (5 mM Tris/HCl, 250 mM Sucrose, 1 mM EDTA, 1 mM EGTA, 1.5 mM MgCl2, 1 mM PMSF, pH 7.4), containing 0.008 % digitonin, and were incubated for 10 min on ice. Cells were centrifuged for 10 min at 15.000 g at 4 °C. The supernatant was collected as cytosolic fraction. The pellet was washed 2 times, resuspended in mitobuffer, and collected as organelle-fraction.

### Electrophoresis, Electrophoretic mobility shift assay, and Western blotting

Cell lysates were thawn on ice and centrifuged at 4 °C for 15 min at 13.000 rpm. The protein content of the supernatants was analyzed according to Bradford (BioRad). 20-30 µg of total protein were diluted in sample buffer (5-fold: 0.3 M Tris/HCl, pH 7, 50 % glycerol, 5 % SDS, 10 mM EDTA, 0.1% bromphenol blue), were reduced with 100 mM DTT for 20 min at RT. Proteins were separated by SDS PAGE using Mini-Protean TGX stain-free 4-20 % precast gels (BioRad).

For the electrophoretic mobility shift assay (EMSA), we synthesized IRD700 labeled RNA used as Hif2α-IRE sequence before GGCUCCUGAGGCGGCCGUACAAUCCUCGGCAGUGUCCUGAGACUGUAUGGUCAGCUCAGCCCAUG ^17^ (Metabion, Germany). According to manufacturer and Anderson et al., the IRE was folded in 60 mM KCl, 6 mM HEPES, 0.2 mM MgCl_2_ via heating to 90°C for 2 minutes and cooling on ice for 5 min. 2 µl of 0.1 µM RNA was mixed with 20 µg cell lysate or 1 µM recombinant IRP1 and 10 µM recombinant Trx1 or 300 mM DTT or 10 mM β-mercaptoethanol in 20 mM HEPES pH 7.4. Samples were incubated with 20 µg/ml BSA for 10 minutes at RT followed by addition of heparin to 0.5 µg/µl. For supershift, 1 µl of 1:20 diluted selfmade anti-IRP1-antibodies ^75^ was added for another 2 minutes. SDS PAGE was run as native gel (native PAGE buffer: 25 mM Tris, 192 mM Glycine, pH 8.3) at 120 V in the dark. Afterwards the probe was detected at 700 nm using Odyssey Infrared Imaging System (LI-COR Bioscience).

The 2D diagonal redox SDS PAGE was performed modifying the protocol described by ^76^. In brief, 40 µg cell lysate was mixed with sample buffer, denatured for 10 min at 96°C before it was separated at a 4-20 % PROTEAN TGX stain-free gel (BioRad) at 200 V for 30 min. The protein lane was cut out and reduced in 250 mM DTT in 1-fold sample buffer at 65 °C. The gel was washed in 1-fold sample buffer, before the proteins were alkylated for 20 min at a reciprocal shaker using 100 mM NEM in 1-fold sample buffer. The gel was washed in 1-fold sample buffer again before it was placed at a 4-20 % PROTEAN TGX IPG gel. A molecular weight marker was added and the gel lane was overlaid with 1% agarose in 1-fold sample buffer. The proteins were separated at 50 V for 10 min, followed by 200 V for 30 min.

The protein content of all gels (and later the blots) was imaged after activation using BioRads stain-free technology with the ChemiDoc XRS+ System. Western blotting was performed using the Trans-Blot Turbo RTA Transfer Kit (BioRad), according to the manufacturers protocol using PVDF membranes. Membranes were blocked with Tris-buffered saline containing 0.05 % Tween 20, 5% nonfat milk powder and 1 % BSA for 1 h at RT and were incubated with specific primary antibodies against Trx1 ^65^, PPAT (kindly supplied by R. Lill, Marburg Germany), IRP1 (abcam ab126595 for 1D gels and for 2D gels see reference ^75^, ferritin (abcam ab75973), and the transferrin receptor (Life technologies 13-6890) at 4 °C overnight. Using horseradish peroxidase (HRP)-coupled secondary antibodies (BioRad) and the enhanced chemiluminescence method antigen–antibody complexes were detected by the ChemiDoc XRS+ System (BioRad). The densitometric analysis was performed using ImageJ and the ImageLab 5.0 Software (BioRad).

### Identification of interaction partners

Co-immunoprecipitation of IRP1 and Trx1 was performed using CnBr activated sepharose. 100 mg sepharose beads were incubated with IRP1 antibodies ^29^ corresponding to the equivalent of 1 ml serum. After washing steps with PBS, the sepharose beads were blocked with 1 M ethanol amine. 20 µl antibody conjugated sepharose beads were incubated with 1 mg protein lysate at 4 °C over night. After removing unbound proteins, the sepharose and bound proteins were denatured for 10 min at 95 °C and Trx1 was stained after performing a reducing SDS PAGE and Western blotting.

The intermediate trapping was performed by immobilizing Trx1C35S on a HisTrap column. Following reduction with 10 mM DTT, 10-20 mg HeLa cell lysate were loaded on the column. Following extensive washing, trapped proteins were eluted with 10 mM DTT, were precipitated over night at 4 °C with 20 % TCA and were centrifuged for 15 min at 13.000 rpm and 4 °C. The pellet was washed with ice cold acetone, centrifuged again and resuspended in lysis buffer. The presence of IRP1 was analyzed by SDS-Page and Western Blotting using the IRP1 antibody described in ^29^ and mass spectrometry. Samples were prepared for mass spectrometric analysis by in-gel reduction, alkylation and digestion essentially as described earlier ^77^. Digested peptides were extracted from the gel, dried and resuspended in 0.1 % trifluoroacetic acid. Peptide samples were subjected to liquid chromatography (Ultimate 3000 rapid separation liquid chromatography system, Thermo Fisher Scientific) coupled tandem mass spectrometry (QExactive plus, Thermo Fisher scientific) as described in ^77^. Briefly, after a 2 h separation of peptides on C18 material, separated peptides were injected via a nano-electrospray source into the mass spectrometer operated in positive, data-dependent mode. First, precursor spectra were recorded at a resolution of 140.000 followed by the quadrupole enabled isolation of up to ten precursors which were subsequently fragmentation by higher-energy C-trap dissociation and resulting spectra recorded at a resolution of 17.500. Peptide and protein identification was carried out with MaxQuant (version 1.6.17.0) on the basis of 75.777 homo sapiens entries (UniProt KB proteome, UP000005640, downloaded on 20210127). The “match between runs”, label-free quantification and iBAQ option were enabled, other parameters were not changed according to the standard settings.

### Molecular modeling and Molecular Dynamics simulation

The suggested conformational changes required for the binding of the FeS cofactor to human Trx1 were modeled using SPDBV ^78,79^. Based on the structure of reduced human Trx1 (pdb entry 1ert), we scanned the loop database for potential alternative conformations of the loop connecting alpha helix 3 and strand 4 of the central beta-sheet. Glu70 and the *cis*-Pro75 were used as anchoring residues. The best fit was obtained with a model loop obtained from structure 2nacA (residues 25-28). The resulting structure was relaxed by energy minimization. The integrity and quality of the structure was confirmed using ProSA-web ^80^, the z-score of -6.9 was within the range of scores typically found for native proteins of this size. The proposed holo-structure of dimeric Trx1 was modeled based on the holo-structures of human Grx2 (2ht9) and human Grx5 (2wul), electrostactic calculations were performed as described in ^52^. Apo-IRP1, *i.e.* the missing loop containing cysteinyl residues 503 and 506, was modeled using Swiss Model (https://swissmodel.expasy.org/) with pdb entry 3sn2 as template. Structures were displayed using UCSF Chimera (https://www.cgl.ucsf.edu/chimera/).

Molecular Dynamics simulations were performed in Gromacs (Version 2021.3) ^81^ with Amber99ff-ILDN force field ^82^. The parameters for the [2Fe-2S] cluster coordinated from Carvalho *et al* ^83^ were used. The model protein was solvated with TI3P water in a cubic box under periodic boundary condition with 1 nm distance from the protein to the edge of the box. The simulation box was neutralised with Na^+^ and Cl^+^ ions.

Prior to the production simulations, the system underwent a three-step equilibration. The first energy minimisation was performed with steepest descent algorithm until the system converged to 1.000 kJ*mol-^1^*nm^-1^. The next equilibration step was performed with NVT ensemble with constant temperature of 300K for 100 ps, and the last step with NPT ensemble with constant pressure of 1bar and temperature of 300K. The time steps of 2 fs were used in the simulations. LINCS ^84^ algorithm was used for constrains of hydrogen bonding interactions. For pressure and temperature coupling the Parrinello-Rahman method ^85^ and modified Berendsen thermostat ^86^ were used, respectively. For the calculation of long-range electrostatic interactions the Particle Mesh Ewald ^87^ method was used, whereas for the short-range (both Coulomb and van-der-Waals) the Verlet cut-off scheme with 1.5 nm cutoff distance was used. The analysis of the simulations were performed with Gromacs-intern tools, UCSF Chimera ^88^ was used for the graphical representation of protein structures.

### Phylogenetic and statistical analysis

Sequences for Trx1 and IRP1/Aco1 were collected from the uniprot.org and pubmed databases. Alignments and the radial phylogenetic tree were created using the CLC Sequence Viewer 8. Alignments were used to analyze the presence of the active site Cys residues as well as the extra Cys73. Statistical significance was calculated using GraphPad Prism 5 (GraphPad Software) when n ≥ 4 using Students t-test (*: p<0.05; **: p<0.01; ***: p<0.001).

## Acknowledgements

The authors wish to thank C.A. Helm and A. Gröning (Greifswald) for help with the differential scanning calorimetry, Elias Arnér (Stockholm, Sweden) for providing mammalian TrxR, Ingrid Span (Düsseldorf/Erlangen, Germany) for help with the anaerobic chamber, Roland Lill (Marburg, Germany) for providing anti-PPAT-antibodies, and Rick Eisenstein (Madison, USA) for helpful discussions. This work was supported by grants from Plan Cancer (to JMM: n° 18CB010-00) and the Deutsche Forschungsgemeinschaft (DFG, German Research Foundation), in particular the Priority Program SPP 1927 (to CB: BE 3259/5-1,2, BE 3259/6-1, 417677437/GRK2578; to CHL: LI 984/3-1,2, LI 984/4-1, GRK1947-A1, to CSM: SCHU 1480/4-1, and to VS: SCHU 1251/17-1,2).

## Author contributions

C.B., E.M.H., M.G., L.T., C.O.S., and C.H.L. performed biochemical and cellular experiments, E.M.H., S.S., Y.B., and O.H. cloned and purified Trx mutants, L.M.J., C.S.M., and V.S. performed Mössbauer spectroscopy, G.P. performed mass spectrometry, R.N. and C.S. synthesized and provided (NH) ^57^Fe(SO) ·6H O, Y.B. and C.H.L. performed bioinformatic analyses including MD simulations, J.M.M. provided critical input and material, C.B., E.M.H., and C.H.L. wrote the manuscript, all authors approved the text.

