## Supplementary Material for "FeS-cluster coordination of vertebrate thioredoxin regulates suppression of hypoxia-induced factor 2α through iron regulatory protein 1"

[1]Department of Neurology, Medical Faculty and University Clinic Düsseldorf, Heinrich Heine University, D-40225, Germany, [2]Institute for Biochemistry and Molecular Biology, University Medicine Greifswald, D-17475, Germany, [3]Department of Otorhinolaryngology, University Hospital Essen, D-45147, Germany, [4]Institute of Molecular Medicine, Proteome Research, Medical Faculty, Heinrich Heine University Düsseldorf, D-40225, Germany, [5]Department of Physics, University of Kaiserslautern-Landau, D-67663, Germany, [6]Institute of Biochemistry, University of Greifswald, D-17487, Germany, [7]Univ. Grenoble Alpes, CEA-IRIG and LBFA, Inserm U1055, Grenoble, F-38000, France

*Carsten Berndt

*Christopher Horst Lillig

**This PDF file includes:**

Figures S1 to S8
Tables S1 to S2
3 References

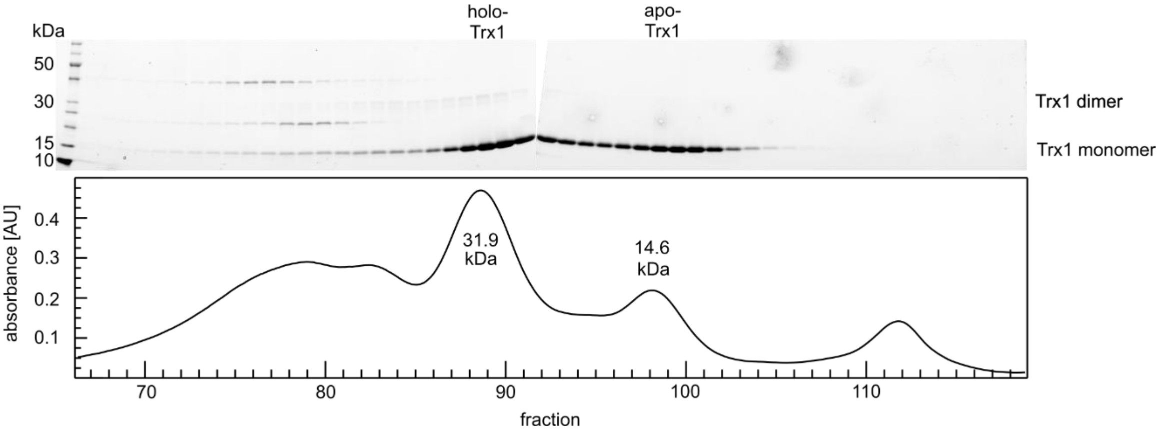


**Fig. S1 Mouse Trx1 forms dimers and monomers after *in vitro* reconstitution and is the main/single protein in the respective gelfiltration peaks**. Reconstituted mTrx1 was applied to gelfiltration. The peaks fit to the theoretical MW of mTrx1 monomer (13.84 kDa) and dimer (27.76 kDa). The corresponding gels demonstrate that purified Trx1 is the only protein in these two peak fractions and thereby the protein leading to the recorded spectra.

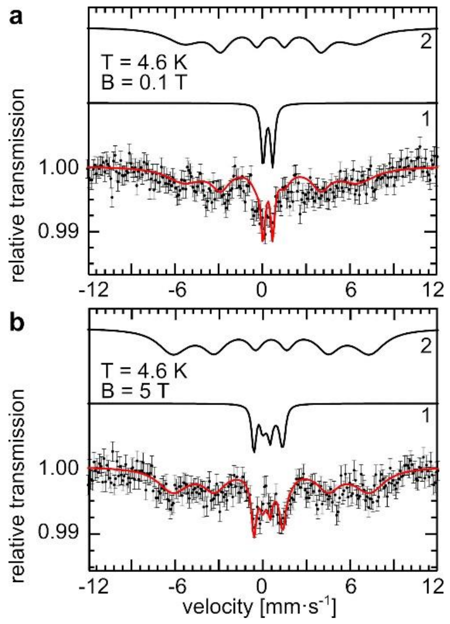


**Fig. S2 Mössbauer spectra of reconstituted human Trx1 obtained at 4.6 K in the presence of external magnetic fields.** a) 0.1 T, b) 5 T. The field direction was parallel to the γ-ray. The simulated spectra of fractions 1 and 2 (see text and Tab. S1) were included on top of the recorded spectra.

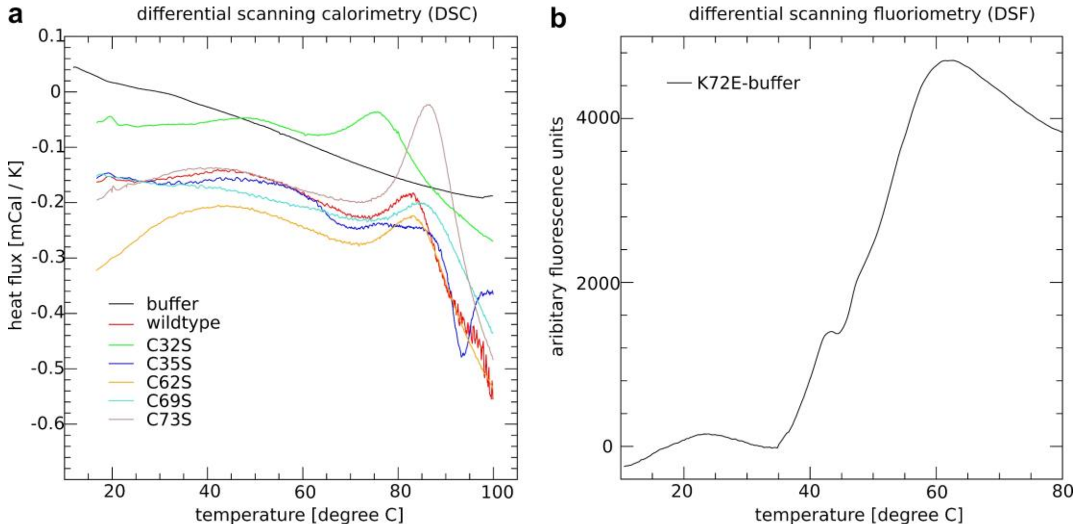


**Fig. S3 Thermal stability of the recombinantly expressed Trx1 mutants.** a) Differential scanning calorimetry: The recombinant proteins at 40 µM in PBS buffer were heated at 1 K per minute from 20 to 100°C. The resulting heat flux was recorded. In all cases, denaturation started above 70°C, for the $T_m$ see peaks. b) Differential scanning fluorimetry: The recombinant protein at 10 µM in PBS buffer was heated at 0.2 K per minute from 20 to 80 °C in the presence of sypro orange. Here, the $T_m$ corresponds to the center of the increasing part of the curve, it was determined by fitting the data to the Boltzmann equation (see Methods section). The thermal stability of wild-type Trx1 determined here was lower compared to previous studies (see for instance [SI1]). This was likely caused by the use of different buffer systems and the use of His-tagged proteins in our study. The calculated denaturation temperatures of the Trx1 mutants are summarized in table S2.

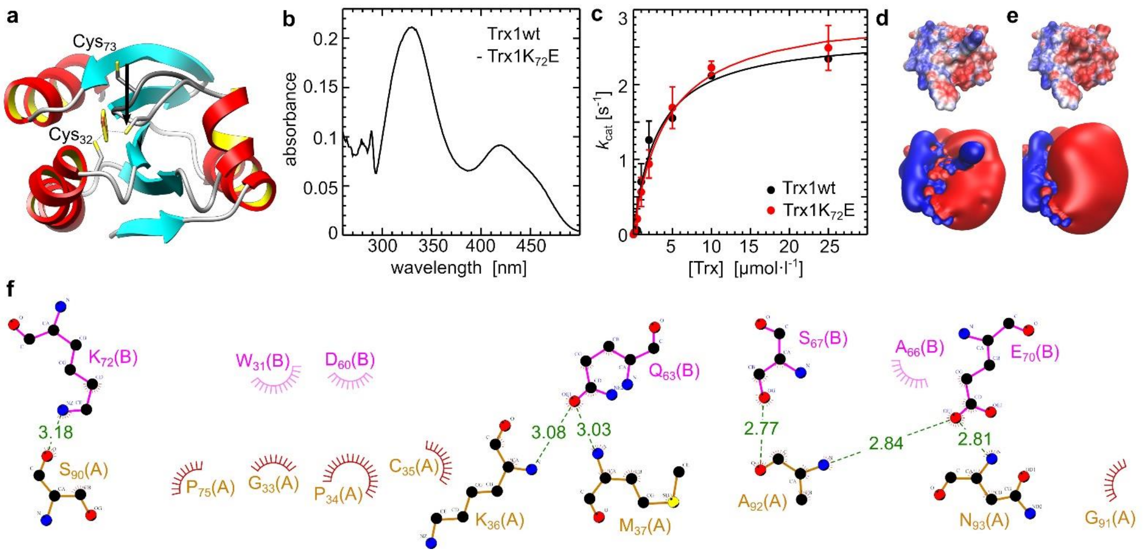


**Fig. S4 Conformational changes in human Trx1 upon binding of the iron cofactor.** a) Conformational changes in the loop containing Cys73 required for FeS cluster ligation (dark gray apo-Trx1, light gray holo-Trx1). The arrow represents app. 6-7 Å. b) Difference UV-Vis spectra between wildtype and K72E Trx1 demonstrates the lack in [2Fe-2S] absorptivity in the mutant. c) Michaelis-Menten plot of TrxR1 activity with Trx1wt and Trx1K72E as substrate in a coupled assay with insulin to keep Trx1 in the oxidized state (n=6, mean ± SD). d-e) Electrostatic surface potential (upper panel) from -100 to +100 mV and elelctrostatic isosurfaces at +/- 25 mV of the K72E mutant (d) and wildtype protein (e). f) DimPlot showing the interaction between the two Trx1 monomers in a dimer (dashed lines: hydrogen bonds, arcs: hydrophopic contacts). The plot is based on a representative frame of the MD simulations.

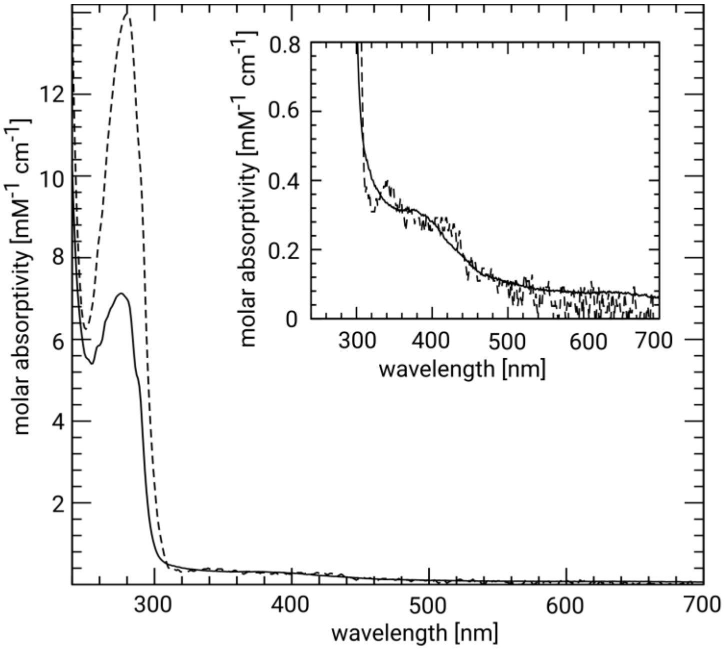


**Fig. S5 *E. coli* Trx1 and human Trx2 are not FeS cluster proteins.** UV-Vis spectra of *E. coli* Trx1 (dashed line) and human Trx2 (straight line). The proteins were expressed and purified in the same way as human and mouse Trx1. *E.c.*Trx1 was cloned into the expression vector pET15b using the restriction sites NdeI and BamHI and the following primers: fwd: 5' CACACACATATGAGCGATAAAATTATTCACCTGACTGACGACAG and rev: 5' CACACAGGATCCTTACGCCAGGTTAGCGTCGAGGAAC. Construction of the mTrx2 plasmid is described in [SI2].

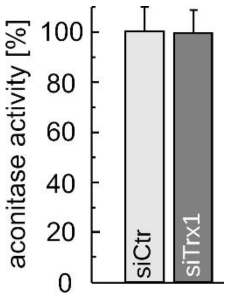


**Fig. S6 Trx1 deficiency does not affect aconitase activity.** Cytosolic aconitase activity of control and Trx1-depleted HeLa cells cultivated under 1 % $O_2$ (n=3, mean ± SD, **: p<0.01, Students T-test).

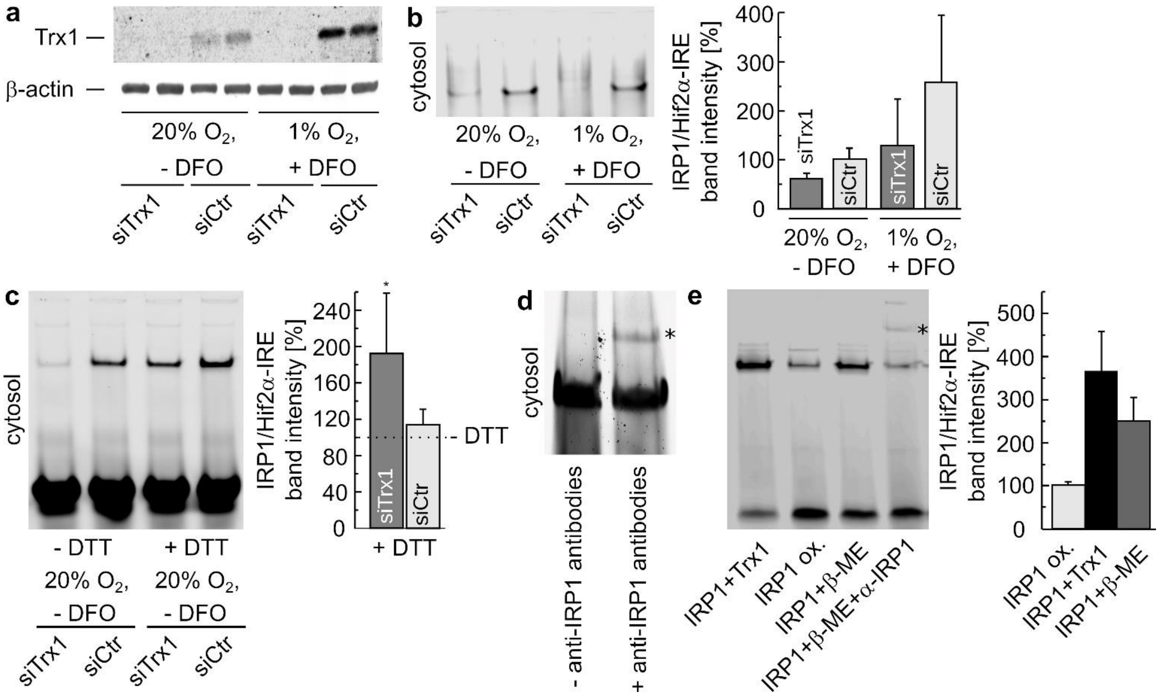


**Fig. S7 Electrophoretic mobility shift assays (EMSAs) using a Hif2α-IRE probe.** a) Successful knock-out of Trx1 via siRNA in the HeLa cells used for the EMSAs. b) EMSAs of cytosolic fractions of HeLa cells ± hypoxic (1 % $O_2$) and iron-limiting (+ iron chelator desferoxamine (DFO)) conditions ± Trx1 depletion (siTrx1). n = 3, mean ± SEM. c) Cytosolic fractions of HeLa cells grown as indicated were treated with or without 300 mM DTT before EMSA. n = 5, mean ± SEM, *: $p<0.05$, Students T-test. d) Formation of a supershift complex (asterisk) consisting of probe, IRP1, and anti-IRP1 antibodies in cytosolic fractions of HeLa cells propagated under 1 % $O_2$ + DFO and electroporated with control siRNA. e) Recombinant IRP1 was incubated ± recombinant apo-Trx1 (10 µM) or ± β-mercaptoethanol (β-ME, 10 mM) with or without presence of anti-IRP1 antibodies for the formation of superhift complexes (asterisk) (n = 3-4, mean ± SEM).

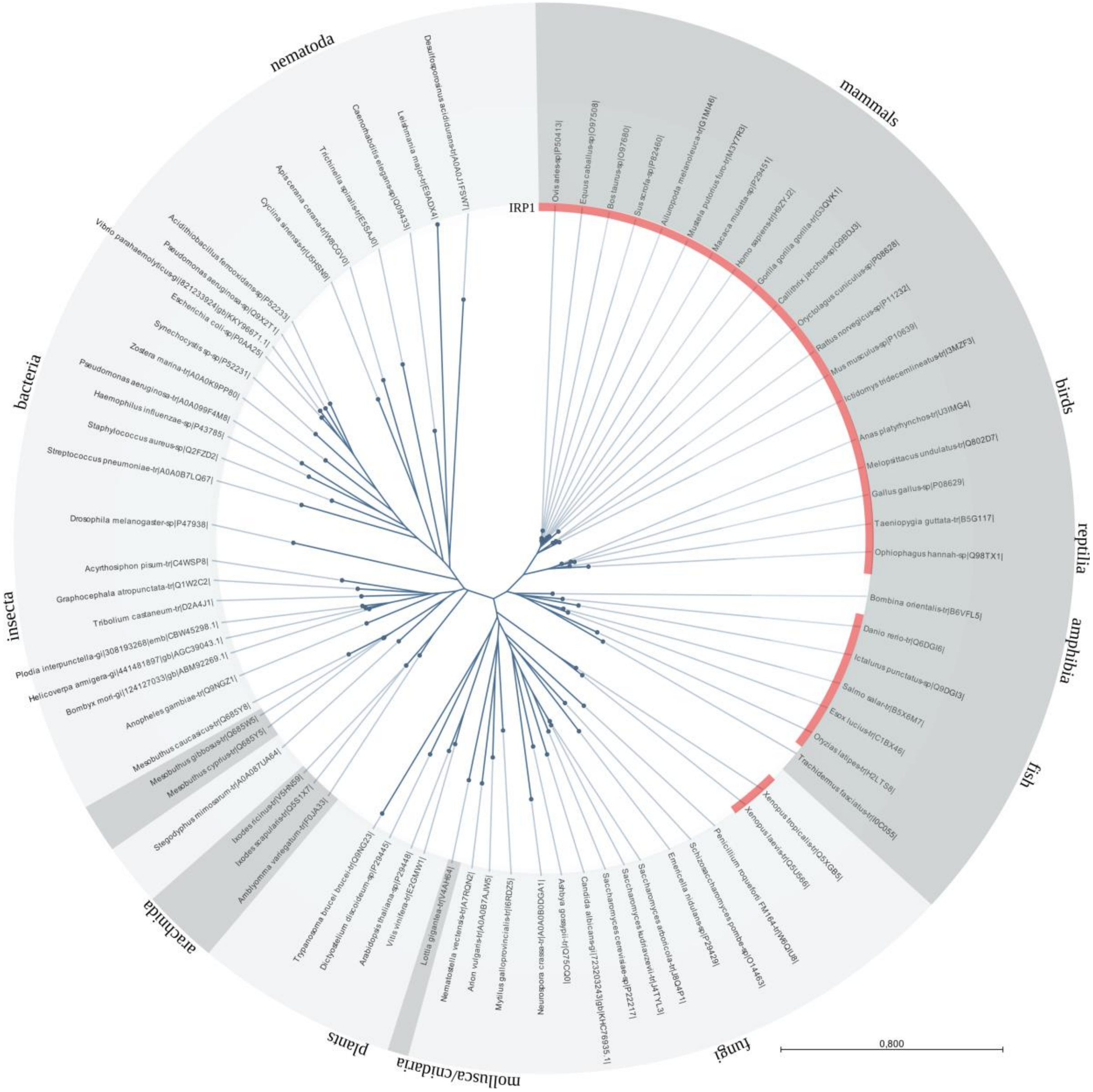


**Fig. S8 Phylogeny of Trx1 homologues.** We have performed sequence comparison between Trxs from various species, classes, and kingdoms. All species contain the active site motif CxxC. The radial phylogenic tree shows the species that contain the extra Cys73 essential for FeS coordination (highlighted in grey); mainly vertebrates, arachnid species and one mollusk. The figure also includes the species in which IRP1 has been identified (depicted in red). Alignments and the radial phylogenetic tree were created using the CLC Sequence Viewer 8.

**Table S1 Mössbauer parameters of reconstituted holo-Trx1 obtained at 4.6 K in presence of external magnetic fields.** Holo-Trx1 was reconstituted *in vitro* and concentrated to 2.5 mM. The protein was frozen in liquid nitrogen and analyzed by Mössbauer spectroscopy as outlined in the experimental procedures, see also Fig. S2. Fraction 1 was simulated by means of the spin-Hamiltionian formalism assuming a diamagnetic (S = 0) ground state [SI3].

| parameter | fraction 1 | fraction 2 |
|---|---|---|
| | S=0 | |
| $\delta$ (mm $s^{-1}$) | 0.35 ± 0.02 | 0.56 ± 0.10 |
| $\Delta E_Q$ (mm $s^{-1}$) | 0.66 ± 0.03 | 0 |
| $\eta$ | 0 ± 0.5 | - |
| $\Gamma_{(5T)}$ (mm $s^{-1}$) | 0.33 ± 0.03 | 2.5/1.9/1.2 ± 0.50 |
| $\Gamma_{(0,1T)}$ (mm $s^{-1}$) | 0.33 ± 0.03 | 3/1.5/1 ± 0.50 |
| $B_{HF(5T)}$ (T) | - | 42 ± 1 |
| $B_{HF(0,1T)}$ (T) | - | 37 ± 1 |
| area (%) | 20 ± 2 | 80 ± 2 |

**Table S2 Denaturation temperatures of the Trx1 mutants.** The temperatures where half of the proteins were found to be denatured was extracted from Fig. S3.

| protein | denaturation temperature °C |
|---|---|
| human Trx1 wildtype | 82.8 |
| human Trx1 C32S | 75.6 |
| human Trx1 C35S | 83.5 |
| human Trx1 C62S | 83.5 |
| human Trx1 C69S | 85.4 |
| human Trx1 C73S | 86.3 |
| human Trx1 K72E | 49.7 |
